# Mapping functional, structural, and transcriptomic correlates of homotopic connectivity

**DOI:** 10.1101/2025.06.10.657943

**Authors:** Annie G. Bryant, James M. Shine, Ben D. Fulcher

## Abstract

Homotopic functional connectivity (HoFC)—the synchronous activity between homologous regions of the left and right cortical hemispheres—is a hallmark of inter-hemispheric brain architecture, yet its biological underpinnings remain incompletely understood. Here, we characterize spatial variation in resting-state HoFC and its relation to dominant natural axes of anatomical, functional, and transcriptomic organization across the human cerebral cortex. We show that the regional distribution of HoFC, from the lowest in association areas (especially limbic) to the highest in primary sensorimotor regions, is preserved in neuropsychiatric disorders in a separate neuroimaging cohort (including schizophrenia, bipolar I disorder, and attention-deficit hyperactivity disorder) despite disease-associated perturbations in magnitude. Regional variation in HoFC is not fully explained by properties of physical embedding or vascular innervation in the brain, suggesting involvement of alternative physiological processes. Furthermore, we find that regions with stronger HoFC are more functionally connected within and across hemispheres at rest, suggesting that homotopic functional coupling is positively associated with brain-wide functional synchrony. Our results collectively suggest that HoFC is inherently aligned with a fundamental natural axis of macroscopic cortical organization and shared global signal—advancing our understanding of the biological correlates of cortical homotopy and suggesting potential mechanisms to investigate in the future.

## 1 Introduction

The left and right hemispheres of the cerebral cortex exhibit a spectrum of lateralized functions and integrated processes that collectively underpin behaviors ranging from sensory perception to higher-order cognition. Coordination between the hemispheres is characterized in part by functional connectivity between ‘homotopic’ regions of the brain—homologous areas in the left and right hemispheres—which comprises a core feature of macroscopic brain organization. Indeed, converging evidence indicates that in the resting-state brain (i.e., in the absence of a cognitive task), homotopic regions show greater synchrony than intra-hemispheric or heterotopic region–region pairs [1–6]. Such inter-hemispheric coupling emerges in embryonic development [7] and is maintained throughout the lifespan, with healthy aging-related increases in sensorimotor areas and decreases in higher-order associative regions [8]. Homotopic functional connectivity (‘HoFC’) has been documented across species—including rodent [9–11], zebra finch [5], and macaque [12, 13]—and is among the most reproducible and temporally robust features observed in resting-state functional magnetic resonance imaging (fMRI) [13, 14], as well as in rodent wide-field calcium imaging [15]. While the pronounced functional coupling between homologous regions is conserved across development and evolution, its relationship with macroscale properties of brain structural and functional organization remains incompletely understood.

Computational modeling analyses have suggested that properties of both geometric embedding (e.g., physical distance or number of streamlines between a pair of regions) and homophilic attraction (e.g., similarity in gene expression or cytoarchitecture) jointly shape the cortical connectome [16]. Within this network-based framework, homotopic connections coincide with greater similarity in gene-expression patterns, receptor density profiles, cortical laminarity, and hemodynamic connectivity than heterotopic connections [2]. HoFC appears to be physically supported by dense structural interconnections via callosal fibers [17], which are considerably more numerous between homotopic than non-homotopic regions in the human brain [18]. Homotopic connections appear somewhat anomalous in terms of cortical geometry, however. For example, Salvador et al. [19] demonstrated that HoFC does not decrease as a function of distance between the two homotopic regions, which has been independently validated in subsequent studies [13, 20]—suggesting that HoFC does not adhere to the general inverse association between resting-state functional connectivity (FC) magnitude and inter-regional distance [19, 21]. Moreover, while the majority of callosal fibers innervate homotopic regions (as opposed to intra-hemispheric or heterotopic region–region pairs) [18], Uddin et al. [22] reported preserved HoFC following complete commissurotomy, in which the corpus callosum is surgically severed in the case of treatment-resistant epilepsy. Since a commissurotomy disrupts monosynaptic callosal projections, the extant HoFC is hypothesized to be supported (in part) through alternate polysynaptic routes—potentially via the anterior commissure, which is known to connect homotopic regions in the temporal cortex [23]. Indeed, O’reilly et al. [24] reported that inter-hemispheric coupling in the rhesus monkey was largely preserved after severing callosal projections, as long as the anterior commissure was left intact. Resting HoFC has been reported in mammals without a corpus callosum by ontogeny [5], further pointing to alternative (or potentially supplementary) physical mechanisms underpinning homotopic connectivity. Collectively, this raises important questions about the neurophysiological mechanisms that underpin inter-hemispheric connectivity throughout the brain.

One of the dominant axes of macroscopic organization in the cortex follows an organized and gradual spatial transition of functional connectivity properties [25, 26]. The first principal gradient of functional connectivity, derived in Margulies et al. [25], places primary sensorimotor regions at one end and higher-order transmodal and association regions at the other [27]—aligning with a canonical ‘functional hierarchy’ in the cerebral cortex [28]. This spatially ordered functional gradient naturally maps onto the topological hierarchy of the cortex [26], originally described to encapsulate synaptic architecture in the primate visual cortex [29, 30]—where ‘topological’ refers to the number of synaptic steps away from the retina. The brain’s topological organization is hypothesized to mechanistically underpin hierarchical functional specialization [31, 32]. Emerging evidence suggests that HoFC also systematically varies along this principal functional gradient; for example, Stark et al. [1] characterized the regional heterogeneity in HoFC, identifying a spectrum ranging from the lowest HoFC (transmodal association regions) up to the highest HoFC (primary sensory-motor regions). The authors posit that the higher linear correlation between BOLD activity in primary sensory regions reflects a ‘default state’ of inter-hemispheric synchrony, which decreases in favor of lateralized hemisphere-specific processing as information is passed up the putative functional hierarchy [25, 26]. Other groups have reported similar patterns from primary unimodal to heteromodal to association cortices [33, 34]—though inter-hemispheric synchrony may ramp up among higher-order regions in the context of a complex task [35, 36].

Amid this regional heterogeneity, we currently lack a comprehensive account of how region-specific variation in HoFC interfaces with broader macroscopic cortical organization—a gap that must be bridged to better understand inter-hemispheric communication in the resting brain, in both health and disease. Here, we focus on regional HoFC variation across the cortex in the resting state, aiming to investigate how the spatial distribution of HoFC relates to macroscale gradients of functional, anatomical, and transcriptomic variation across the cortex. Hypothesizing that resting HoFC is linked to a region’s position along the dominant axes of functional and anatomical organization in the cortex, respectively, we specifically compared regional HoFC with brain-wide maps capturing anatomical hierarchy, resting-state functional connectivity, and transcriptomic variation. We also characterize the robustness of regional variation in HoFC magnitudes in the context of distinct neuropsychiatric disorders (in an independent neuroimaging cohort), where disruptions to inter-hemispheric coupling are frequently reported [37–46], and examine how homotopic coupling relates to cortex-wide resting-state functional connectivity within the ipsilateral and contralateral cortices. We also explore several potential physiological bases for cortical HoFC derived from physical embedding, structural connections, and vascular innervation. Collectively, our analyses contribute to our understanding of the structural, functional, and molecular correlates of homotopic connectivity in the human cerebral cortex, suggesting promising avenues for future hypothesis-driven investigation.

## 2 Results

In order to quantitatively investigate macroscale patterns of inter-hemispheric coupling, we characterize the spatial distribution of homotopic functional connectivity (HoFC) across the human cerebral cortex. We analyze openly available data from the Human Connectome Project (HCP) [47], with preprocessed group-consensus structural and functional connectomes provided by the ENIGMA Consortium [48] and described in depth previously [49]. Briefly, Larivière et al. [48] first computed individual structural and functional connectomes for each of *N* = 207 participants; for the functional connectome, negative connections were set to zero and Fisher’s *r*-to-*z* transformation [50] was applied (as in previous graph-theoretical analyses [51, 52]), before averaging across participants to yield a consensus connectome. More details about this dataset are included in the Methods (Sec. 5.1.1). We first assess how the spatial variation in HoFC magnitude relates to maps of different properties of macroscopic cortical organization along anatomical, functional, and transcriptomic gradients included in the *neuromaps* library [53] (in Sec. 2.1). After characterizing these relationships, we then evaluate the robustness of the spatial variation in HoFC magnitudes to three neuropsychiatric disorders in which inter-hemispheric synchrony is reportedly disrupted, querying the UCLA Consortium for Neuropsychiatric Phenomics dataset [54] (in Sec. 2.2). Finally, we interpret HoFC in the context of the broader topological landscape of the cerebral cortex by comparing the average functional network strength with HoFC magnitude (in Sec. 2.3).

### 2.1 Homotopic functional connectivity varies along a fundamental axis of cortical organization

We first characterized the spatial distribution of HoFC magnitudes across the cerebral cortex from a consensus connectome derived from *N* = 207 individuals from the HCP S1200 release (see Methods, Sec. 5.1.1). For each cortical region in the parcellation atlas from Desikan et al. [55] (34 regions per hemisphere), we examined the *z*-scored Pearson correlation coefficient (*R*_Z_) between the left and right hemispheres from the group consensus connectome. This approach yielded a cortex-wide map of HoFC magnitudes, allowing us to quantitatively compare regional HoFC gradients with other properties of macroscale cortical organization. As shown in brain maps in Fig. 1A, we observed a clear gradient of HoFC magnitudes across the cortex, from the lowest values in the entorhinal cortex (*R*_Z_ = 0.05) and temporal pole (*R*_Z_ = 0.03; not visible in Fig. 1A) to the highest in the superior parietal cortex (*R*_Z_ = 0.71) and lateral occipital cortex (*R*_Z_ = 0.70). This pattern is consistent with previous analyses using the HCP dataset [33, 34] as well as different imaging cohorts [1, 20, 33], supporting prior findings that resting-state HoFC is highest in primary motor, visual, and somatosensory regions and lowest in association cortices.

**Figure 1.**
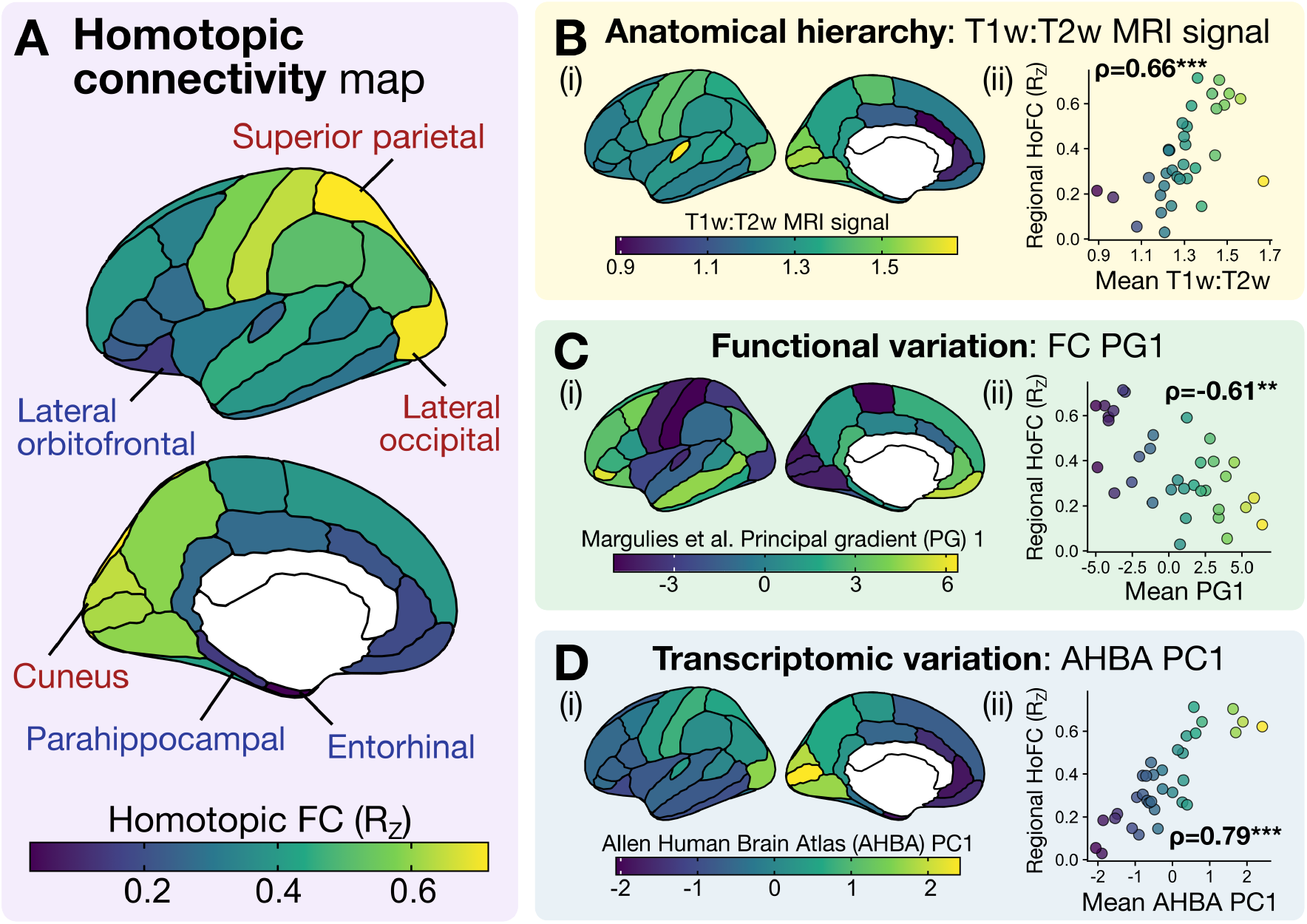
Regional homotopic functional connectivity (HoFC) variation aligns with a fundamental axis of macroscale cortical organization. **A**. The group consensus *z*-transformed Pearson correlation coefficient (*R*_Z_) between homotopic region pairs is depicted on the cortical surface. Regions with the highest HoFC are annotated with red text (superior parietal, lateral occipital, and cuneus) and regions with the lowest HoFC visible from medial/lateral views are annotated with blue text (lateral orbitofrontal, parahippocampal, entorhinal). **B**. (i) The mean T1w:T2w MRI signal is plotted for each region (averaged between the two hemispheres) in the same parcellation atlas (from Desikan et al. [55]) as the HoFC maps in **A**. (ii) The mean T1w:T2w MRI signal (*x*-axis) is compared with the HoFC value (*y*-axis) per region, with the Spearman rank correlation (*ρ*) annotated. **C**. (i) The mean first principal component of gene expression from the Allen Human Brain Atlas (AHBA PC1) is plotted for each region (averaged between the two hemispheres). (ii) As in **B**(ii), the AHBA PC1 value (*x*-axis) is compared with the HoFC value (*y*-axis) per region. **D**. (i) The mean first principal gradient (PG1) of functional connectivity (FC) from Margulies et al. [25] is plotted for each region (averaged between the two hemispheres). (ii) As in **B**(ii), the PG1 value (*x*-axis) is compared with the HoFC value (*y*-axis) per region. ^***^, *P <* 0.001; ^**^, *P <* 0.01; single-hemisphere spin test with 10,000 permutations [56]

Margulies et al. [25] demonstrated that the first principal gradient (PG1) of resting-state functional connectivity systematically varies across the cortical sheet in a way that tracks the topological hierarchy of laminar projections, ranging from primary sensory regions to transmodal to association cortices [26]. Here, we focus on properties of concerted variation in macroscale cortical organization and their relation to a ‘fundamental axis’ of the cerebral cortex [28], with ‘hierarchy’ referring specifically to the anatomical organization of laminar feed-forward and feed-back patterns of inter-areal anatomical connections (originally identified in non-human primate cortex with histological tract-tracing) analysis [30, 57, 58]. Specifically, we examine three distinct types of graded spatial variation across the cortical sheet using curated reference maps from the `neuromaps` toolbox [53]. Given the proposed links between HoFC and white matter microstructural properties [3], we first queried the ratio of T1-weighted to T2-weighted (T1w:T2w) signal from structural MRI, which is sensitive to intracortical myelin content (as well as several other microstructural properties of neurons) [59]. Burt et al. [60] subsequently demonstrated that T1w:T2w is a reliable non-invasive proxy for the anatomical hierarchy of the cortex. We additionally queried the first principal gradient (PG1) of functional connectivity (FC), which captures maximal variance in resting-state functional connectivity across the cortical sheet, positioning primary sensorimotor regions at one end and higher-order association regions at the other [25]. Lastly, we examined transcriptional organization across the cortex using the first principal component (PC1) of brain-wide gene expression from the Allen Human Brain Atlas, (AHBA) [61], which captures maximal variance in gene expression and provides a proxy for cell-type distributions across the cortex [60, 62, 63]. We compared the regional HoFC variation (as shown in Fig. 1A) with the mean value in the corresponding area of each map of macroscale cortical organization, averaging between the left and right hemispheres for each in the latter case. Statistical significance was evaluated using a non-parametric single-hemisphere spin test [56] with 10,000 permutations (see Methods for details, Sec. 5.3.1). Of note, the T1w:T2w and AHBA gradients exhibit a positive correlation in their respective spatial variations [60] and a negative correlation with that of PG1 magnitudes; in other words, T1w:T2w and AHBA values are generally negatively correlated with the anatomical hierarchy of the cortex, while PG1 values exhibit a positive association.

As depicted in Figs 1B-D, we found that the spatial variation in HoFC magnitude is significantly correlated with all three maps of macroscale cortical organization—in a manner that suggests HoFC is inversely associated with the anatomical hierarchy in the cortex. Specifically, we observed a positive correlation to the T1w:T2w map (*ρ* = 0.66; *P*_spin_ = 5 *×* 10^−4^, single-hemisphere spin test with 10,000 permutations) and transcriptomic AHBA PC1 map (*ρ* = 0.79; *P*_spin_ = 2 *×* 10^−4^) and a negative correlation to the PG1 map (*ρ* = −0.61; *P*_spin_ = 1 *×* 10^−3^). Regions with the highest-magnitude resting HoFC (*R*_Z_ ≥ 0.6) exhibited among the highest values of T1w:T2w MRI signal (Fig. 1B), corresponding to lower positions along the anatomical hierarchy. These same high-magnitude HoFC regions also sit at the apex of AHBA PC1 gene expression values (Fig. 1D), which tracks a molecular gradient in which higher values correspond to greater microglial and CA1 pyramidal neuron density and downregulation of cell signaling and modification pathways [64]. Similar alignment between T1w:T2w and AHBA PC1 gradients was reported in Burt et al. [60], suggesting these two properties converge toward a common axis of anatomical and transcriptomic regional variation. Lastly, the high-magnitude HoFC regions exhibited among the lowest functional PG1 values (Fig. 1C), which maps to the primary unimodal end (rather than association) of the dominant axis of functional connectivity described in Margulies et al. [25]. Collectively, our results suggest that HoFC spatial variation tracks primary axes of anatomical hierarchy and macroscale organization across the cortical sheet.

### 2.2 Spatial variation in homotopic functional connectivity is preserved across neuropsychiatric disorders

We next posed two follow-up questions: (1) How reproducible is the spatial variation in resting-state HoFC in another young and clinically normative population?; and (2) Is this macroscale HoFC gradient altered in the context of disease? For the latter, we focused on neuropsychiatric disorders, which are generally attributed to dysfunction across distributed brain networks [65–68], with widespread alterations to functional coupling throughout the cerebral cortex. Mounting evidence indicates that HoFC is reduced across the brain in a multitude of neuropsychiatric disorders, including anxiety and depression [37–39], autism spectrum disorder [40–42], bipolar disorder [43], and schizophrenia [44–46], suggesting that reduced HoFC may serve as a transdiagnostic biomarker [69]. However, it remains to be clarified whether this general reduction in HoFC exhibits characteristic spatial changes in a given disorder or manifests diffusely throughout the cortex. To address both of these questions, we investigated the spatial distribution of resting-state HoFC magnitudes in participants in a neurotypical control group (*N* = 116) or those diagnosed with one of three neuropsychiatric disorders included in the UCLA Consortium for Neuropsychiatric Phenomics (CNP) study [54]: schizophrenia (SCZ; *N* = 48), bipolar I disorder (BP; *N* = 49), and attention-deficit hyperactivity disorder (ADHD; *N* = 39). More information about data acquisition and preprocessing with this cohort is provided in the Methods (Sec. 5.1.2).

First, comparing the HoFC map from neurotypical controls in the UCLA CNP cohort (cf. Fig. 2A) to that of the HCP cohort (cf. Fig. 1A) demonstrated general concordance between the two maps, with a Spearman correlation of *ρ* = 0.62 (*P* = 5 *×* 10^−4^, single-hemisphere spin test with 10,000 permutations [56]). The direct spatial comparison is shown in more detail in Fig. S1, with overall visual agreement in the pattern of higher HoFC in primary sensorimotor regions and lower HoFC in higher-order transmodal and association areas. Next, having demonstrated consistent spatial variation in HoFC magnitudes in this independent control cohort, we investigated the spatial variation in HoFC in each of the three disorders. As shown in Figs 2B(i)-D(i), the distribution of HoFC magnitudes across the cortex was visually similar between each neuropsychiatric disorder cohort (averaged across participants) and that of the control group—although magnitudes were visually lower in SCZ in particular. To directly examine the nature of HoFC changes across the cortical sheet in these three disorders, we computed the mean difference in HoFC *R* values between each disorder and the control cohort per region, yielding a Δ*R* = *R*_Disorder_ − *R*_Control_ value for each region. These Δ*R* values are plotted on the cortical surface in Figs 2B(ii)-D(ii), revealing that most regions exhibited reduced HoFC to some degree relative to controls, consistent with previous reports [43–46]. The most pronounced HoFC differences appeared in SCZ, in which three cortical areas showed a change of Δ*R <* −0.1: pars opercularis (Δ*R* = −0.13), middle temporal gyrus (Δ*R* = −0.12), and superior temporal gyrus (Δ*R* = −0.11). By contrast, most regions in BP and ADHD showed smaller differences in HoFC (−0.05 ≤ Δ*R* ≤ 0.05), though the banks of the superior temporal sulcus (‘bankssts’) exhibited more pronounced HoFC reduction in BP with Δ*R* = −0.11. Overall, the Δ*R* distributions reflect heterogeneous and relatively low-magnitude HoFC perturbations in each disorder.

**Figure 2.**
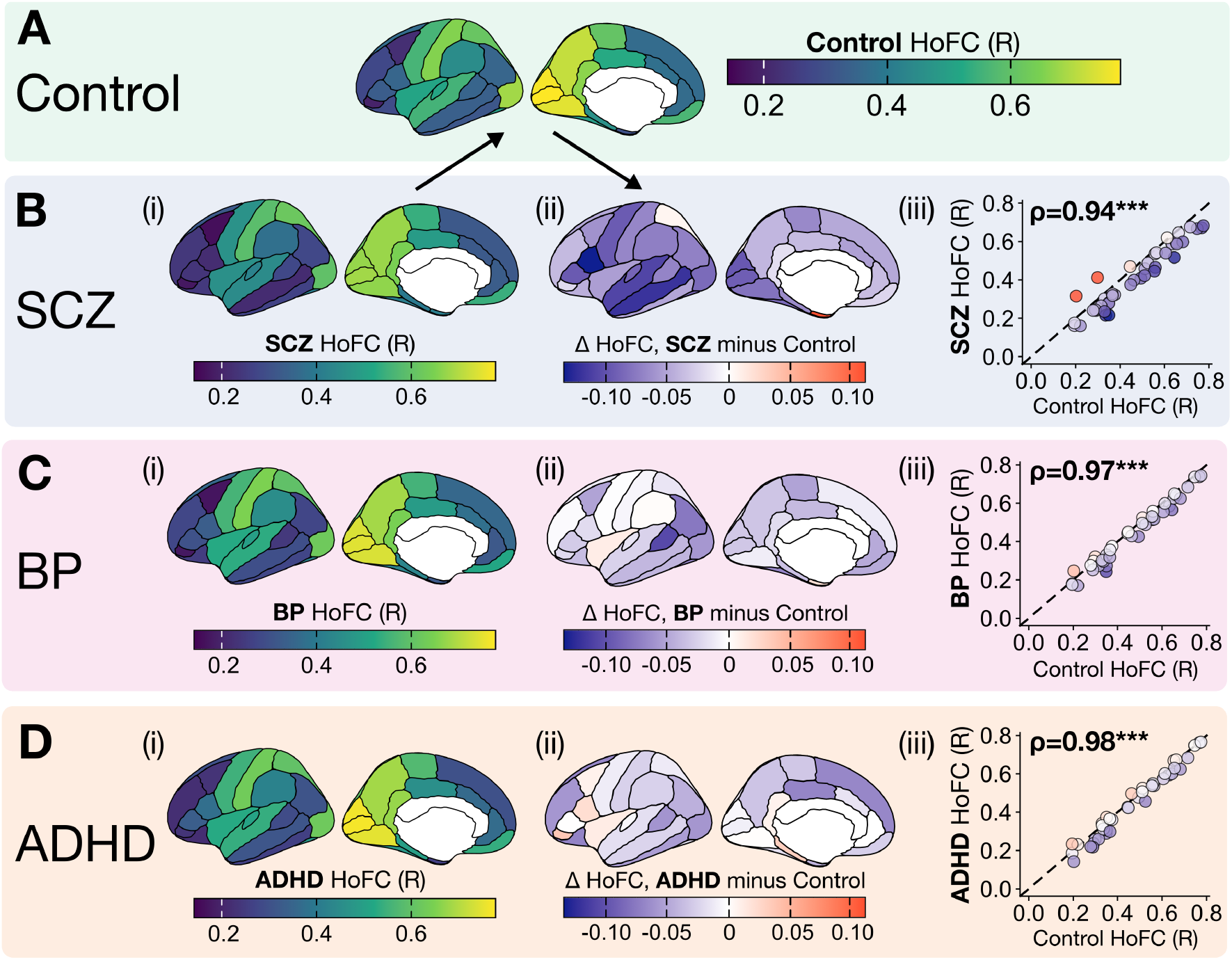
The macroscale HoFC regional gradient is preserved across neuropsychiatric disorders, despite relatively homogenous global decreases in inter-hemispheric coupling. **A**. The regional HoFC gradient is depicted on the cortical surface for control participants (N=116) in the UCLA CNP cohort. HoFC values reflect group-averaged Pearson correlation coefficient values (*R*); note that correlation values were not *z*-scored in the preprocessing pipeline for this dataset. **B**. (i) The regional HoFC gradient is depicted on the cortical surface for schizophrenia (SCZ) participants (N=48) in the UCLA CNP cohort. (ii) For each brain region, the difference in HoFC (Δ HoFC, in units of *R*) between SCZ versus control participants is plotted on the cortical surface. (iii) The regional HoFC values are plotted for control (*x*-axis) versus SCZ (*y*-axis) participants, with the Spearman rank correlation (*ρ*) indicated in the top left corner. Points reflect brain regions (total of 34) and are colored as in (ii), and the identity line with slope=1 is shown as a black dashed line. **C**. For bipolar I disorder (BP, N=49), the regional HoFC gradient (i), difference relative to controls (ii), and scatter plot compared with control values (iii) are plotted as with SCZ in **B. D**. For attention-deficit hyperactivity disorder (ADHD, N=38), the regional HoFC gradient (i), difference relative to controls (ii), and scatter plot compared with control values (iii) are plotted as with SCZ in **B**. ^***^, *P <* 0.001; single-hemisphere spin test with 10,000 permutations [56]

Against the backdrop of generally reduced HoFC (to varying degrees) in SCZ, BP, and ADHD, we next quantified the extent to which the HoFC spatial gradient observed in the neurotypical control cortex is altered in each of these neuropsychiatric disorders. In Figs 2B(iii)-D(iii), we compared the HoFC (*R*) per cortical region in each diagnostic group (*x*-axis) versus that of the UCLA CNP control cohort (*y*-axis, from Fig. 2A). The spatial gradient of HoFC was largely preserved in all three disorders, with Spearman’s *ρ* = 0.93 in SCZ, *ρ* = 0.97 in BP, and *ρ* = 0.98 in ADHD. While most regions fell below the identity line (dashed black in Fig. 2D), meaning Δ*R <* 0, computing the line of best fit per disorder (not shown) yielded slopes between *m* = 0.91 (SCZ) and *m* = 1.02 (ADHD)—suggesting that despite heterogeneous disease-associated alterations across the cortical sheet, the overall regional variation in HoFC appears to be robust and reproducible across cohorts.

### 2.3 Homotopic connectivity is linked to broader resting-state functional network topology

While prior work has demonstrated that homotopic region pairs exhibit overall greater linear correlation in BOLD activity than intra-hemispheric or heterotopic connections [2–6, 13], to our knowledge, the relationship between HoFC and general properties of functional connectivity in the resting brain has not been investigated previously. To address this gap, we sought to clarify how regional HoFC magnitude relates to average node strength in the functional connectome, both within and across hemispheres. We aggregated the connections per region–region pair into a single group FC matrix, with intra-hemispheric connections in one matrix triangle and inter-hemispheric connections in the other, as depicted schematically in Fig. 3A. The resulting group-averaged functional connectome is plotted in Fig. 3B, which visually demonstrates a striking level of symmetry between intra-hemispheric functional connections (left triangle) and inter-hemispheric connections (right triangle) across all region–region pairs. Regions are ordered according to their mean HoFC magnitude, revealing a spectrum ranging from overall higher FC values (both within and across hemispheres) in regions with higher HoFC to lower cortex-wide FC values in regions with lower HoFC. For each region, we computed its overall average (non-homotopic) nodal strength within the intra- and inter-hemispheric functional networks, defined as the average functional connectivity (in *R*_Z_) between each region and each of the other 33 regions in the atlas from Desikan et al. [55]. As shown in Fig. 3C, the average strength per region was strongly correlated with HoFC magnitudes, both within (*ρ* = 0.86; *P*_spin_ = 1 *×* 10^−4^, single-hemisphere spin test with 10,000 permutations) and across hemispheres (*ρ* = 0.92, *P*_spin_ = 1 *×* 10^−4^). This indicates that the macroscale gradient in resting-state HoFC also mirrors that of average FC strength generally, both within and across hemispheres.

**Figure 3.**
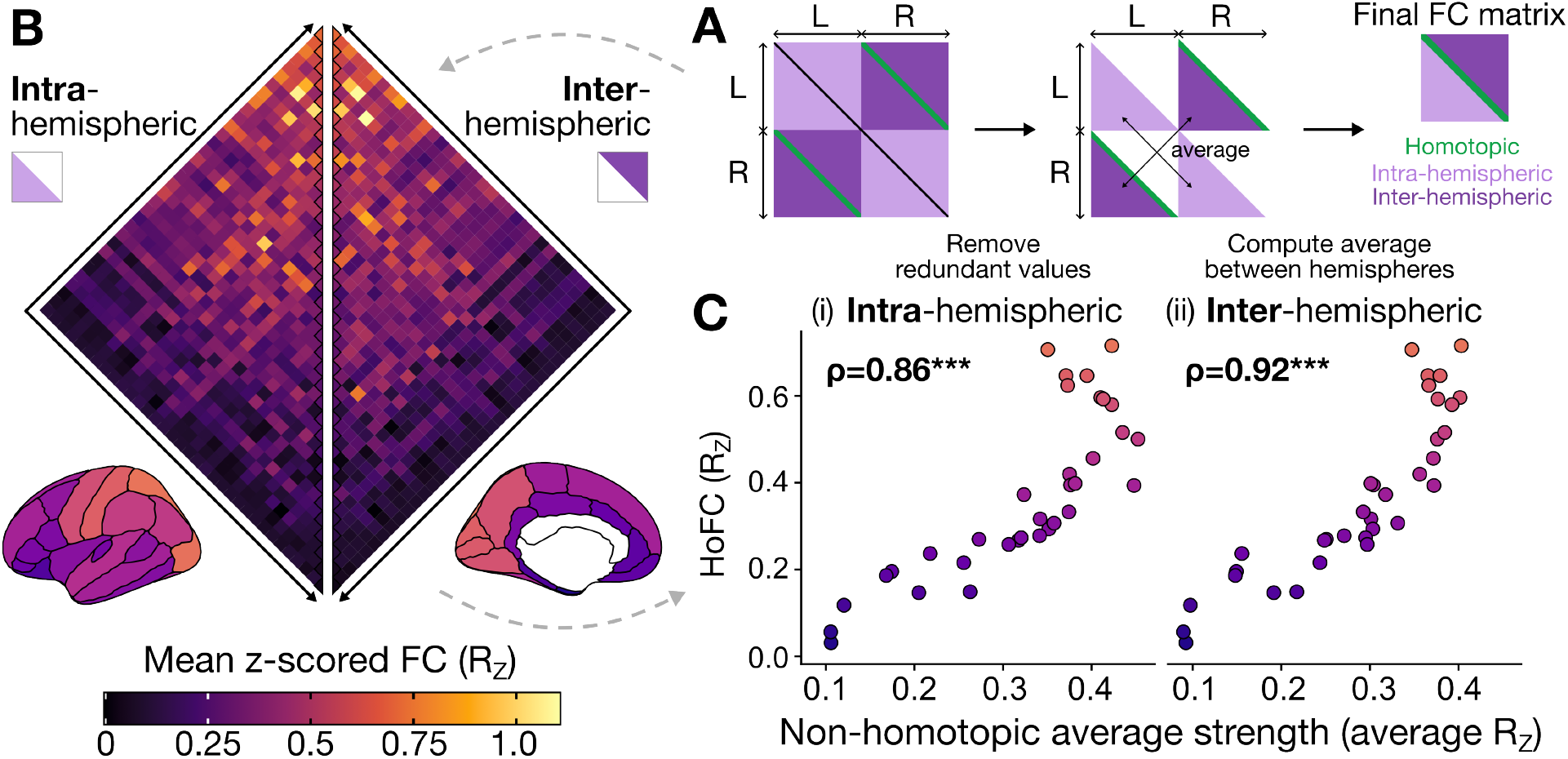
Homotopic functional connectivity (HoFC) is linked to regional degree within and between hemispheres in the functional connectome. **A**. Our approach to compare hemisphere-averaged connections within versus across hemispheres is depicted schematically. Starting with the full 68*×*68 functional connectome (with the atlas from Desikan et al. [55]), we first divided the connectome into four quadrants (L ↔ L, L ↔ R, R ↔ L, and R ↔ R). Within each quadrant, correlation values above and below the diagonal line are symmetric (and therefore, redundant), so only the lower four triangles were retained. To generalize across left and right hemispheres, we then computed the average intra-hemispheric connections (by averaging over L ↔ L and R ↔ R per region–region pair) and average inter-hemispheric connections (by averaging over R ↔ L and L ↔ R per region–region pair). This yielded one final matrix, with homotopic connections along the diagonal, and one triangle for intra- and inter-hemispheric functional connections, respectively. **B**. The resulting group-averaged functional connectome (hemisphere-averaged from the process depicted in **A**) is shown, with values representing the mean *z*-scored Pearson correlation across all individuals in the HCP cohort from ENIGMA (see Methods, Sec. 5.1.1). The left triangle depicts intra-hemispheric connectivity, while the right triangle depicts inter-hemispheric connectivity. The nodes along the center line indicate homotopic connections, and the same HoFC values shown in the heatmap are also plotted on the cortical surface below to guide visual interpretation with the same color scale. **C**. For each cortical region, the strength is computed as the average FC value (*R*_Z_) to all other regions within the (i) ipsilateral hemisphere or (ii) contralateral hemisphere and plotted on the *x*-axis versus the HoFC value for the corresponding region on the *y*-axis. In each case (intra- or inter-hemispheric), the Spearman rank correlation (*ρ*) is indicated at the top left corner. ^***^, *P <* 0.001; single-hemisphere spin test with 10,000 permutations [56]

### 2.4 Homotopic functional connectivity is spatially related to variation in vascular innervation, but not structural architecture

In order to better understand plausible candidates for the physiological underpinnings of HoFC in health and disease, we next investigated properties related to structural connectivity, physical embedding, and vascular pathology. Structural connectivity within the mammalian cortex follows an exponential distance rule, such that the probability of a physical connection between two areas and the strength of the connection both decrease exponentially with increasing physical distance [71, 72]; this effect has been replicated in human structural connectivity analysis [73, 74]. Resting-state FC generally exhibits a similar spatial constraint [19, 75, 76], and the structure–function coupling magnitude varies along the anatomical hierarchy of the cortex [77–79]. Cortico-cortical HoFC strength is also reportedly linked to structural connectivity integrity between region–region pairs [3, 80]—as in resting-state FC in general [21]. However, prior work also indicates that HoFC does not decrease with inter-region distance, with greater connectivity between homotopic regions than would be expected based on their physical distance [13, 19, 20]—perhaps due to the specificity of cross-hemisphere connectivity. In light of these findings, it has yet to be clarified whether HoFC is associated with cortical variation in structural architecture. To test this, we compared the resting HoFC gradient in Fig. 1A with: (1) anatomical connectivity, quantified as the log-transformed number of streamlines estimated using diffusion tensor imaging; and (2) spatial proximity, quantified as the Euclidean distance between the centroid vertex of each region on the cortical surface. The group-consensus structural connectome analyzed here is from the same participants as with the functional connectome, all preprocessed in Larivière et al. [48]; more details are provided in the Methods (Sec. 5.2.3).

As shown in Fig. 4B, the cortical gradient of resting HoFC was not significantly correlated with structural connectivity, estimated as the log_10_-transformed streamline count (*ρ* = 0.12, *P*_spin_ = 0.5, single-hemisphere spin test with 10,000 permutations [56]). Of note, the preprocessed structural connectome was generated with a distance-dependent consensus threshold approach [81] to retain edges reliably detected across the *N* = 207 participants while preserving individual-level connection length distributions. In the resulting structural connectome provided in Larivière et al. [48], for 12 of the 34 regions in the atlas from Desikan et al. [55], homotopic anatomical connections were not reliably detected across participants after adjusting for streamline length, such that the estimated streamline count was approximated as zero after log-transforming all other streamline counts (corresponding to the 12 regions plotted along the zero line on the *x*-axis in Fig. 4B). We confirmed in a robustness analysis that in the absence of thresholding, with a subset of 100 structural connectomes (out of the *N* = 207 in the ENIGMA-provided dataset [48]) preprocessed with a similar pipeline [82], there was no significant association between anatomical connectivity and HoFC across the cortex (*ρ* = 0.18, *P*_spin_ = 0.31; Fig. S2). Similarly, there was no significant association between spatial proximity and HoFC magnitude (*ρ* = 0.16, *P*_spin_ = 0.3), as shown in Fig. 4C. Some of the closest regions along the cortical midline (separated by ≤20 mm) showed relatively low HoFC (*R*_Z_ ≤ 0.4); in contrast to regions like the postcentral gyrus, which exhibited among the highest HoFC (*R*_Z_ = 0.64), despite the left and right regions sitting 85.3 mm apart on average (in the fsaverage template surface). These findings indicate that properties of anatomical wiring and physical embedding do not account for regional variation in HoFC strength—suggesting that the spatial HoFC gradient may be related to additional factors that are (at least partly) dissociable from direct structural architecture.

**Figure 4.**
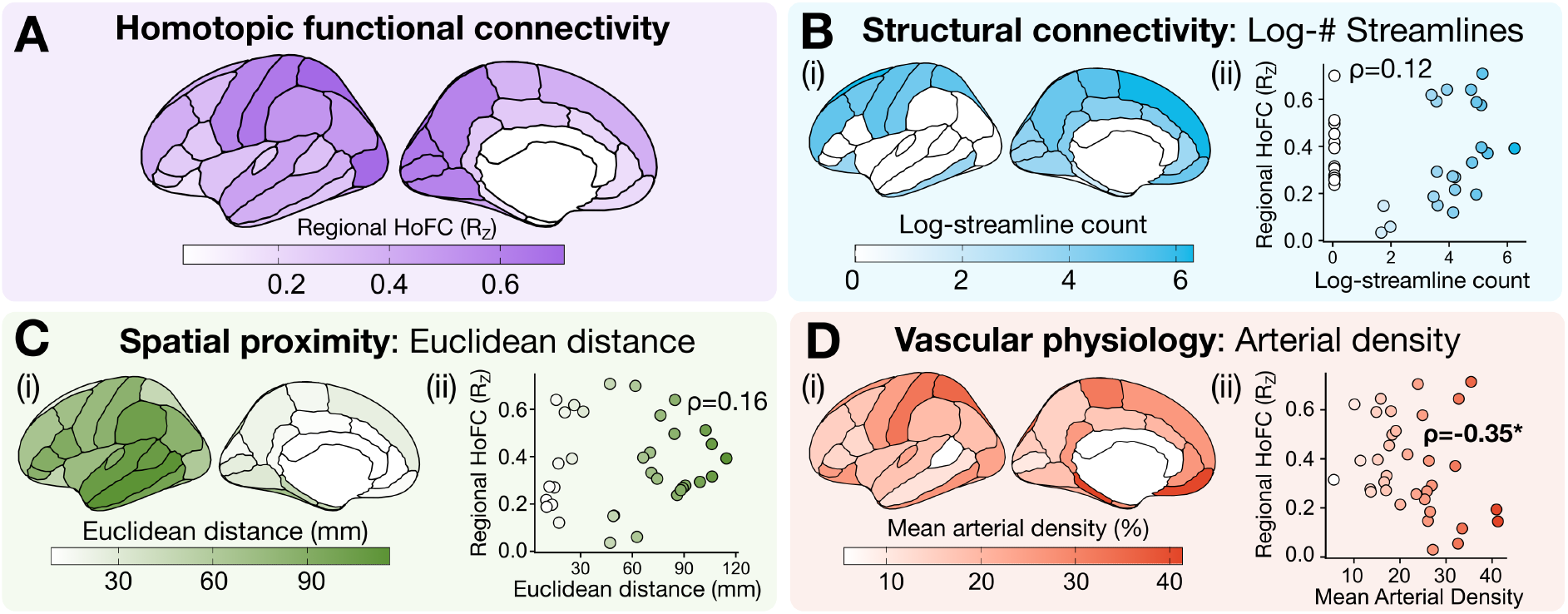
Regional HoFC is not statistically associated with structural connectivity or Euclidean distance, but it is inversely associated with arterial density. **A**. The same HoFC gradient from Fig. 1A is replicated here to guide visual interpretation, with values projected onto the cortical surface. **B**. (i) The log_10_-transformed streamline count between homotopic region pairs is depicted on the cortical surface. As part of the distance-dependent threshold procedure applied to this group-consensus structural connectome in Larivière et al. [48], region–region pairs which did not reliably exhibit anatomical projections were thresholded out, and the log-count set to zero. (ii) The mean log-streamline count connecting each homotopic region pair (*x*-axis) is plotted against the HoFC for each corresponding region (*y*-axis), with the Spearman rank correlation (*ρ*) annotated. **C**. (i) The Euclidean distance between the centroid vertices of each homotopic region pair is depicted on the cortical surface. (ii) As in **B**(ii), the mean Euclidean distance between each homotopic region pair (*x*-axis) is plotted against the HoFC for each corresponding region (*y*-axis). **D**. (i) The mean arterial density in each region [70] is plotted on the cortical surface, averaged between the two hemispheres per region. (ii) As in **B**(ii), the mean hemisphere-averaged arterial density (*x*-axis) is plotted against the HoFC for each corresponding region (*y*-axis). ^*^, *P*_spin_ *<* 0.05; single-hemisphere spin test with 10,000 permutations [56].

As another possible physiological mechanism that might contribute to homotopic BOLD connectivity, we also examined properties of the cerebral vasculature. BOLD fMRI intrinsically captures neurovascular coupling by measuring the ratio of deoxygenated to oxygenated hemoglobin over time in each voxel [83]. Prior work indicates that homotopic regions exhibit highly synchronous oscillations in vessel diameter [84], and Drew et al. [85] posit that vascular geometry serves as a physiological substrate for HoFC in addition to inter-hemispheric anatomical projections. Moreover, emerging work suggests that asymmetries in capillary transit time heterogeneity [86] and/or venous draining [87] are associated with attenuated BOLD fMRI connectivity between homotopic regions. Given these findings linking vascular innervation properties with HoFC, we reasoned that spatial HoFC variation may relate to differences in the type and density of vascular innervation across the cortical surface. To investigate this, we queried the probabilistic atlases for arterial and venous density in Bernier et al. [70] in the same parcellation space. Arteries carry oxygenated hemoglobin from the heart into the brain; they are generally 1-5 mm in diameter and dilate within a few hundred milliseconds of neural activity onset [88]. By contrast, veins carry de-oxygenated hemoglobin from the brain back into the peripheral vasculature; veins are generally similar in diameter to arteries, though they exhibit dilations much smaller in volume and on a slower timescale than arteries (i.e., tens of seconds after activity onset) [83, 89].

As shown in Figs 4D(i,ii), arterial density was mostly concentrated around medial temporal and frontal regions as well as the lateral parietal cortex, and exhibited a significant negative association with the regional variation in HoFC magnitude (*ρ* = −0.35, *P*_spin_ = 7 *×* 10^−3^, single-hemisphere spin test with 10,000 permutations). This negative association suggests that regions with greater arterial supply tend to exhibit lower HoFC magnitude, which is consistent with emerging work demonstrating that proximity to the nearest artery is inversely associated with a region’s weighted degree centrality (i.e., strength) within the functional connectome [90]. By contrast, venous density was concentrated around visual and inferior frontal/temporal cortices and was not associated with HoFC (*ρ* = −0.08, *P*_spin_ = 0.8; Fig. S3). Collectively, the significant (albeit, relatively low-magnitude) negative correlation between arterial density and HoFC magnitude suggests that cerebral vascularization may be related to inter-hemispheric functional connectivity.

## 3 Discussion

In this study, we provided a comprehensive account of how resting-state homotopic functional connectivity (HoFC)—a key feature of inter-hemispheric organization—varies across the human cortex and relates to structural, functional, and transcriptomic properties. We have shown that regional resting-state HoFC is not spatially homogeneous, but rather exhibits a characteristic regional variation that, in turn, aligns with the dominant axis of anatomical hierarchy and macroscale functional and transcriptomic organization across the cortical sheet (Fig. 1). This spatial gradient was robust to heterogeneous disease-associated perturbations in region-level HoFC (in a separate neuroimaging cohort [54]), most of which corresponded to relatively low-magnitude decreases across the cortex (Fig. 2). Regions that were more functionally connected within and across hemispheres in the resting state also exhibited greater homotopic coupling, suggesting that HoFC and overall functional connectivity strength co-vary along a similar gradient. Furthermore, we demonstrated that regional HoFC variation in the cortex cannot be fully explained by structural architecture or vascular factors (Fig. 4), suggesting that other physiological mechanisms may be supporting resting-state HoFC.

Our findings are consistent with previous reports of inverse associations between HoFC and position along a fundamental axis [28] of macroscale cortical organization, from primary sensorimotor to association regions [1, 33, 34], though the specific mechanisms mediating this relationship will need to be directly tested in the future. For example, prior work has established that primary sensory/motor cortices are more heavily myelinated [91] and exhibit higher T1w:T2w signal ratios [92]. The T1w:T2w ratio is sensitive to multiple microstructural properties of the gray matter [59, 93] and is a hypothesized (inverse) correlate of the anatomical hierarchy of laminar differentiation across the cortex [60]. We observed a significant positive association between HoFC and T1w:T2w gradients across the cerebral cortex, suggesting that HoFC decreases with ascending position along the putative anatomical hierarchy. Of note, Shafiei et al. [94] reported that the gradient of T1w:T2w values—along with that of AHBA PC1 values—is strongly associated with the dominant axis of resting-state neurophysiological fMRI dynamics across the cerebral cortex. Collectively, our findings are congruent with the ‘default state’ of inter-hemispheric synchrony proposed by Stark et al. [1], with the highest HoFC observed in primary sensorimotor regions. The tighter bilateral coupling between these primary sensorimotor regions may suggest greater resting-state symmetry in support of sensory perception and motor coordination, potentially via shared inputs from thalamic drive [95].

The preserved resting-state HoFC gradient in a separate neurotypical cohort and in three neuropsychiatric disorder groups (SCZ, BP, and ADHD) is surprising, given the inter-individual variability in resting-state functional connectivity [96], which is particularly pronounced in neuropsychiatric disorders [97]. This suggests that while most regions exhibited decreased HoFC with heterogeneous magnitudes across the three disorders relative to neurotypical controls, such disease-related perturbations were outweighed by the dominance of the overall HoFC spatial gradient. In a recent meta-analysis, Yao and Kendrick [69] reported that HoFC changes in ADHD were too small and/or variable to be reliably included, instead identifying common HoFC reductions in visual and motor regions in SCZ along with autism spectrum disorder, anxiety, depression, and addiction (n.b., BP was not included in that specific cross-disorder analysis). Prior work in SCZ populations has demonstrated that HoFC reductions are associated with auditory verbal hallucinations [98], social withdrawal [46], and cognitive deficits [99]. To our knowledge, this is the first analysis to demonstrate that the macroscale HoFC gradient is robust to subtle disease-associated alterations at the regional level. Of note, the magnitude of HoFC reductions was most pronounced in SCZ, followed by BP, and least pronounced in ADHD—which is consistent with previous findings that resting-state fMRI dynamics were more distinctively altered in SCZ and BP than in ADHD in this cohort [100]. However, future investigation in an external dataset(s) will be needed to more fully explore this preliminary finding in the context of key covariates, including symptom severity and treatment usage.

Having identified a robust gradient of HoFC across the cortex across multiple participant groups and imaging cohorts, we further established that HoFC magnitude is strongly related to the overall regional FC strength, averaged across all of each region’s functional connections. While the general distribution of brain-wide homotopic connectivity strengths has been compared to that of heterotopic connections [2, 13], the relationship between HoFC in a given region and its other intra- and inter-hemispheric functional connections has not previously been characterized, to our knowledge. Collectively, our functional network analysis suggests that HoFC is higher in regions that are more functionally connected to the rest of the cortex (i.e., higher node average strength; cf. Fig 3). In other words, regions that show greater overall functional connectivity within the ipsilateral hemisphere, and are therefore more functionally central, also exhibit functional synchrony with the contralateral hemisphere, indicative of a more globally shared signal. Our findings suggest that both HoFC and shared global signal are closely related, with decreasing dominance of both measures in the resting-state brain with ascending position along the anatomical hierarchy and the first principal gradient of functional connectivity.

We investigated potential contributions of vascular innervation and/or structural architecture to the resting-state HoFC gradient, finding that only arterial density was (negatively) associated with HoFC magnitude. It is difficult to disentangle biological versus methodological implications in the context of BOLD fMRI, which is itself sensitive to neurovascular coupling [83] and may be less suited to detect arterial rather than venous dynamics [101]. One potential biological interpretation could be that higher arterial density coincides with greater metabolic lateralization, which would in turn reduce the need for synchronous bilateral activity [102–104]—though further hypothesis-driven investigation in an external dataset is warranted to clarify the meaning of the inverse spatial association between HoFC magnitude and regional arterial density. The finding that resting-state HoFC magnitude was not statistically associated with structural connectivity (quantified as log-transformed number of streamlines, estimated from diffusion-weighted imaging) is surprising, given the proportion of callosal fibers innervating homotopic region pairs [18, 105] and previous findings that cortical HoFC is affected by structural connectivity integrity [3, 80]. As with functional connectivity in general, the relationship between structure and function is nuanced and highly variable across both spatial and temporal scales; for example, Shen et al. [13] reported that the number of tract-traced fibers generally correlated with temporal stability of a given homotopic connection, but this pattern was subverted by primary sensory regions (including V1) that do not exhibit direct structural connections. While the quantity of innervating white-matter tracts did not spatially correlate with HoFC magnitude in this analysis, it is worth noting that primary sensorimotor cortices are interconnected via fibers with thicker myelin (i.e., faster conductance) while heteromodal and association areas are interconnected via fibers with thinner myelin (i.e., slower conductance) [105]. It may be that resting-state HoFC is more related to qualities of the structural connections, such as fiber diameter and conductance speed, rather than the overall quantity of fibers [1, 13]; indeed, Mollink et al. [3] reported that HoFC is intrinsically associated with properties of white-matter myeloarchitecture [3].

When interpreting the findings in this paper, one should consider potential limitations that collectively suggest avenues for future investigation. First, while we analyzed two open-access resting-state fMRI datasets— which have been investigated extensively in previous work (HCP and UCLA Consortium for Neuropsychiatric Phenomics)—future replication with an external dataset(s) is needed to validate the biological findings we presented here. Moreover, these datasets were collected with different acquisition parameters and preprocessed using different pipelines, and while the agreement between the two was significant (*ρ* = 0.62), the dependence of HoFC spatial variation on preprocessing methods should be examined. For both BOLD fMRI datasets, we used the atlas from Desikan et al. [55], which is defined based on anatomical boundaries and may not be optimally suited for representing aggregate functional activity [106]. While this atlas was chosen given consistency in nomenclature and topographical position of regions between the left and right hemispheres relative to the other parcellations included as part of the ENIGMA preprocessed dataset [48] (all based on atlases from Schaefer et al. [107]), future investigation with parcellation atlases tailored specifically for homotopic analysis is warranted [108, 109]. As with any associative brain-map analysis, we also point out that the resting-state fMRI data were obtained from different individuals compared to the T1w:T2w, FC PG1, AHBA PC1, and vascular density maps examined herein. Moreover, we exclusively focused on the Pearson correlation coefficient as an index of HoFC, which is specifically sensitive to linear, synchronous coupling between regions. While the Pearson correlation coefficient is empirically well-suited to the spatial and temporal scales of resting-state BOLD fMRI [100], future work should examine the robustness of the cortical HoFC gradient in different imaging modalities (e.g., magnetoencephalography [110]) and with other measures that can capture different types of coupling, such as time-lagged and/or frequency-based connectivity [100, 111]. The present analysis also focused exclusively on resting-state connectivity, though future work is needed to clarify how cortical HoFC evolves, and potentially reorganizes, with increasing task complexity in healthy participants as well as in the context of neuropsychiatric disorders [13, 17, 104, 112].

## 4 Conclusions

In this work, we focus on three aspects of homotopic functional architecture that, to the best of our knowledge, have not been addressed previously: (1) How does regional variation in homotopic connectivity interface with macroscale gradients of cortical organization that include anatomical hierarchy, functional connectivity, and vascular innervation?

(2) Is the characteristic spatial variation in homotopic connectivity robust between cohorts and clinical groups, particularly neuropsychiatric disorders? (3) Does homotopic connectivity relate to a region’s overall strength in the brain-wide functional connectome? Our analyses collectively indicate that: (1) the regional gradient of homotopic connectivity is significantly associated with a fundamental natural axis of macroscale cortical organization, along with arterial density, though not with properties of physical embedding or structural connectivity; and that (2) this spatial variation is remarkably similar in an independent neuroimaging cohort that includes SCZ, BP, and ADHD cases, despite heterogeneous disease-associated perturbations. Finally, (3) homotopic connectivity magnitude is strongly associated with a region’s overall intra- and inter-hemispheric average strength in the functional connectome at rest. If confirmed in future studies that incorporate independent validation datasets, these findings provide important insights into the functional architecture of inter-hemispheric connectivity in health and disease.

## 5 Methods

### 5.1 Neuroimaging datasets

#### 5.1.1 Resting-state fMRI dataset 1: Human Connectome Project 3T fMRI

The first dataset fMRI dataset included in this study is comprised of *N* = 207 unrelated participants (*N* = 83 males, 28.7 *±* 3.7 years of age) from the S1200 release of the Human Connectome Project [47], as curated and preprocessed by the ENIGMA Consortium [48]. We used preprocessed group-averaged functional connectomes included in the ENIGMA Toolbox [48], as in previous work [49, 113]. Details on both imaging acquisition and preprocessing are described in Larivière et al. [48] and are briefly summarized as follows. Resting-state volumes in this dataset were acquired using a gradient-echo, echo planar image (EPI) sequence with the following parameters: TR/TE = 720/33.1 ms, slice thickness = 2.0 mm, 72 slices, 2.0 mm isotropic voxels, frames per run = 1,200. All resting-state fMRI data were preprocessed using the ‘minimal preprocessing’ HCP pipeline [114], which includes corrections for distortion, head motion, and magnetic field bias; skull removal; and intensity normalization. Noise removal was performed using the FIX software to regress out head motion, white matter signal, cardiac pulsation, and signal from arterial and large veins. Corrected fMRI volumes were mapped to MNI152 space and subsequently projected to the cortical surface using a cortical ribbon-constrained mapping algorithm. Normative FC matrices were generated for each participant by computing the pairwise Pearson product-moment correlations (*R*) between resting-state time series of all cortical regions, setting negative correlation values to zero. To generate a group consensus FC matrix, Fisher’s *R*-to-*Z* transformation was applied as in previous work [20, 33] and then averaged across all N=207 participants, such that all FC values from this dataset are presented as *R*_Z_. We queried the ‘aparc’ parcellation (corresponding to the 68-region parcellation using the atlas from Desikan et al. [55]) using the load_fc function to obtain cortical *R*_Z_ values.

#### 5.1.2 Resting-state fMRI dataset 2: UCLA Consortium for Neuropsychiatric Phenomics

In order to characterize regional variation in HoFC in the context of neuropsychiatric disorders, we additionally included resting-state fMRI data from the open-access University of California at Los Angeles (UCLA) Consortium for Neuropsychiatric Phenomics (CNP) LA5c Study [54]. This included cognitively healthy control participants as well as participants diagnosed with schizophrenia (SCZ), attention-deficit hyperactivity disorder (ADHD), and bipolar I disorder (BP). Details of diagnostic criteria and behavioral symptoms have been described previously, along with imaging acquisition details [54]. Resting-state volumes in this dataset were acquired using a T2^*^-weighted EPI sequence with the following parameters: TR/TE = 2s/30 ms, slice thickness = 4.0 mm, 34 slices, 4.0 mm isotropic voxels, frames per run = 152. This resting-state fMRI dataset was preprocessed in previous work [100, 115] as briefly described in the following. Imaging data were preprocessed using the *fmriprep* v1.1.1 software [116] and the independent component analysis-based automatic removal of motion artifacts (ICA–AROMA) pipeline [117]. Additional noise correction was performed to regress out signal from white matter, cerebrospinal fluid, and global gray matter using the ICA–AROMA + 2P + GMR method [115, 118, 119]. Quality control analysis based on noise regression and head motion excluded eight participants from this resting-state fMRI dataset (N=4 control, N=3 SCZ, N=1 ADHD). After preprocessing and quality control, resting-state fMRI data were retained for downstream analysis from a total of N=252 participants: N=116 controls, N=48 SCZ, N=49 BP, and N=39 ADHD. More information about these quality control measures and the demographic composition of the final participant cohort is described in Bryant et al. [100]. The 68-region parcellation from Desikan et al. [55] was applied to extract the noise-corrected 152-length time series per region (corresponding to 304 s), averaging across all voxels per region. FC matrices were generated for each participant by computing the pairwise Pearson product-moment correlations (*R*) between resting-state time series of all cortical region–region pairs. We note that raw Pearson correlation coefficient values (*R*) were retained, without Fisher’s transformation or thresholding, for this dataset. Within each diagnostic group, FC matrices were averaged across participants to yield one group-average FC matrix per group (i.e., one FC matrix each for control, SCZ, BP, and ADHD groups).

### 5.2 Examining brain maps of macroscale functional and molecular organization

#### 5.2.1 Brain maps of anatomical hierarchy, transcriptomic variation, and functional organization

To quantify multiple facets of macroscale organization across the cortical sheet, we examined three brain maps from the neuromaps [53] library: T1w:T2w MRI signal (as a proxy for anatomical hierarchy), AHBA PC1 (as a proxy for cell type density and key transcriptomic variation), and FC PG1 (which orients regions along a principal axis of functional connectivity, from unimodal sensorimotor to higher-order association areas). All map information is provided in the `neuromaps` documentation (https://netneurolab.github.io/neuromaps/listofmaps.html) and briefly summarized in the following. As a proxy of anatomical hierarchy [60], we examined the *hcps1200-myelinmap-fsLR-32k* map of T1-weighted to T2-weighted (T1w:T2w) signal, derived from structural MRI obtained from 417 unrelated participants from the HCP S1200 release [47, 59]. We additionally queried the *abagen-genepc1-fsaverage-10k* map capturing the first principal component (PC) of gene expression in the Allen Human Brain Atlas [61], as computed in Markello et al. [120] to capture the axis of maximal spatial variation in the cortical transcriptome. Lastly, we queried the *margulies2016-fcgradient01-fsLR-32k* map, which quantifies the first principal gradient (PG1) of functional connectivity (FC) from Margulies et al. [25]. Each map was parcellated into the 68-region atlas from Desikan et al. [55] using the `Parcellator` class method, and the average value was computed between hemispheres for each homotopic region pair (e.g., for the cuneus T1w:T2w ratio, the average was computed between the left and right cuneus).

#### 5.2.2 Structural architecture brain maps

We examined two aspects of structural architecture in the cortex: anatomical connectivity and Euclidean distance between homotopic region–region pairs. For anatomical connectivity, we used the group-averaged structural connectome from the same N=207 HCP participants as the fMRI dataset, preprocessed and included in the ENIGMA Toolbox [48]. Details on both diffusion-weighted imaging (DWI) acquisition and preprocessing are described in Larivière et al. [48], and are briefly summarized as follows. DWI volumes were acquired using a spin-echo EPI sequence with the following parameters: TR/TE = 5.52s s/89.5 ms, 1.25 mm isotropic voxels, *b*-values=1,000/2,000/3,000 s/mm^2^, 270 diffusion directions, 18 b0 images. As part of the HCP ‘minimal preprocessing’ pipeline [114], b0 intensity normalization was applied along with correction for susceptibility distortion, eddy currents, and head motion. Structural connectivity matrices were generated per participant using *MRtrix* [121] and whole-brain streamlines were reconstructed using the SIFT2 (spherical-deconvolution informed filtering of tractograms) pipeline [122]. A group consensus structural connectome was derived using a distance-dependent thresholding procedure [81] with log_10_ transformation to reduce variance, yielding a structural connectivity matrix with log-transformed number of streamlines (i.e., fiber density). Streamline counts for region–region pairs in which streamlines were not reliably detected across individuals (after adjusting for length) were set to zero after log-transforming the other streamline counts. We queried the ‘aparc’ parcellation (corresponding to the 68-region parcellation using the atlas from Desikan et al. [55]) using the `load_sc` function. For our robustness analysis presented in Fig. S2, we analyzed diffusion-weighted MRI data from a subset (N=100) of the individuals in the main analysis, as preprocessed separately (though with a very similar pipeline) and described in Fallon et al. [82]. After reconstructing individual-level structural connectomes, the mean streamline count per edge (i.e., region–region pair) was computed, without any thresholding. As with the main analysis, we applied a log_10_ transformation to reduce variance, yielding an unthresholded group structural connectivity matrix with log-transformed number of streamlines. Streamline counts of zero, or less than 1 (i.e., negative after log_10_ transformation), were set to zero after log-transforming the other streamline counts.

To compute the physical distance between each homotopic region–region pair, we identified the centroid vertex (from the pial surface) of each cortical region in the *fsaverage* template space from FreeSurfer. We then calculated the Euclidean distance between the centroid vertices in the left and right hemisphere per region, respectively, using the `cdist` function from the *scipy* package (version 1.13.1) [123] in Python. Since coordinates in the pial surface files per hemisphere (`lh.pial` and `rh.pial`, respectively) are represented in millimeters (mm), the resulting Euclidean distances reflect the mm between each homotopic region–region pair.

#### 5.2.3 Vascular innervation brain maps

We examined two aspects of vascular innervation in the cortex: arterial density and venous density. For both properties, we queried brain maps published in Bernier et al. [70], where the details of imaging acquisition and preprocessing can be found. Briefly, arteries were imaged using time-of-flight angiography (ToF) and veins with susceptibility weighted imaging (SWI), respectively, in a cohort of N=42 healthy individuals (aged 20-31). Arterial and venous density maps were derived from the ToF and SWI maps using a multi-scale Frangi diffusive filter (MSFDF) pipeline [124] which includes denoising, vessel voxel classification, and multiscale vessel enhancement. Vessel centerlines and diameters were extracted and nonlinearly aligned to MNI T1 (0.5mm) space. In order to minimize contamination of the venous tree by arterial signal in SWI, any voxels overlapping with those identified in ToF volumes were excluded. Vascular density maps were computed for gray matter, white matter, and cerebrospinal fluid (CSF) based on voxel-wise vessel segmentation. This yielded voxelwise maps of arterial and venous density, with values ranging from 1-100 that indicate the vessel probability in the corresponding voxel. The 68-region volumetric atlas from Desikan et al. [55] was applied to these voxelwise maps to extract the average arterial and venous density per region, respectively. The average value was computed between hemispheres for each homotopic region pair (e.g., for the cuneus arterial density, the average density was computed between the left and right cuneus).

### 5.3 Statistical analyses

#### 5.3.1 Brain map comparison with homotopic connectivity gradient

We obtained regional homotopic functional connectivity (HoFC) estimates from the group-averaged functional connectomes from the HCP and UCLA CNP datasets as the correlation between the left and right homologs of each region. To compare the spatial variation in HoFC with different brain maps of macroscopic cortical organization, we computed the Spearman rank correlation (*ρ*) with the `spearmanr` function from the *scipy* package [123], comparing across a total of 34 regions. In order to evaluate the statistical significance of correlation estimates while accounting for spatial autocorrelation along the cortical sheet, we performed a single-hemisphere spin test with 10,000 permutations using the method of Alexander-Bloch et al. [56]. This method, implemented with the `nulls.alexander_bloch` function in *neuromaps*, rotates one map (e.g., the T1w:T2w map) along the spherical surface of a single hemisphere, which preserves spatial contiguity while disrupting the spatial alignment between the two compared maps. For each permutation, the Spearman correlation is computed between the permuted and original map (here, the HoFC gradient is always the original map), which collectively yields a null distribution. The resulting *p*-value reflects the proportion of permutations (out of 10,000) with null *ρ* values equal to or greater in magnitude than the observed empirical correlation, *ρ*.

#### 5.3.2 Functional network topology quantification

We examined regional HoFC variation within large-scale functional network topology based on the symmetric 68-region cortical FC matrix provided by the ENIGMA consortium. Left and right hemisphere region labels were collapsed to form a common ‘base region’ name (e.g., consolidating ‘lh-entorhinal’ and ‘rh-entorhinal’ down to ‘entorhinal’). Each connection was labeled as ‘homotopic’ (same region across hemispheres), ‘intra-hemispheric’ (different regions within the same hemisphere), or ‘inter-hemispheric’ (different regions in different hemispheres; sometimes referred to as ‘heterotopic’) based on anatomical pairing. As depicted schematically in Fig. 5.3.2A, we distilled the original group FC matrix down to a 34*×*34 matrix in which homotopic connections occupy the diagonal, intra-hemispheric connections populate the lower triangle, and inter-hemispheric connections comprise the upper triangle. Regions were ordered in both rows and columns according to their HoFC magnitude, ranked from highest to lowest. In order to summarize the average node strength, we computed the mean of the *z*-scored FC values (*R*_*Z*_) between a given region and all 33 other regions in the same hemisphere (for intra-hemispheric connectivity) or the opposite hemisphere (for inter-hemispheric connectivity).

### 5.4 Data visualization

All cortical brain maps were created using the *ggseg* package (version 1.6.6) in R [125]. Scatterplots and heatmaps were generated using the *ggplot2* package [126] (version 3.5.1) and violin plots with the *see* package [127] (version 0.11.0), both in R.

## 5.5 Code and data availability

All code needed to replicate the analyses and visualizations presented in this paper is openly available on GitHub at https://github.com/DynamicsAndNeuralSystems/Homotopic_FC_HCP/. All data included in this paper is openly available for researchers to access. Preprocesed functional and structural connectivity matrices (derived from HCP data, which is openly available [47]) are included as part of the ENIGMA toolbox at https://enigma-toolbox.readthedocs.io/en/latest/pages/05.HCP/ [48]. Resting-state fMRI data from the UCLA CNP study are available for download at OpenNeuro with accession number ds000030 (https://openfmri.org/dataset/ds000030/). All brain maps are included as part of the *neuromaps* Python package unless otherwise stated; for example, the vascular innervation maps [70] are available for download from the Braincharter GitHub repository at https://github.com/braincharter/vasculature/releases/tag/Atlas_v1.0.

## 6 Acknowledgments

We thank A/Prof Joseph T. Lizier, Joshua M. Tan, Dr Giulia Baracchini, Dr Jayson Jenganathan, and Dr Isabella Orlando for insightful discussions that were very helpful in shaping this project. We also thank the Human Connectome Project and UCLA CNP Project, and their respective participants, for openly sharing this excellent data resource. We also thank the ENIGMA Consortium for openly sharing the preprocessed structural and functional HCP connectomes as part of their toolbox, as well as the Network Neuroscience Lab at the Montreal Neurological Institute for sharing the *neuromaps* toolbox and the many curated brain-maps contained therein. High-performance computing facilities provided by the School of Physics at The University of Sydney supported analyses and results presented here.

## Supporting Information for ‘Mapping functional, cytoarchitectonic, and transcriptomic underpinnings of homotopic connectivity’

**Figure S1.**
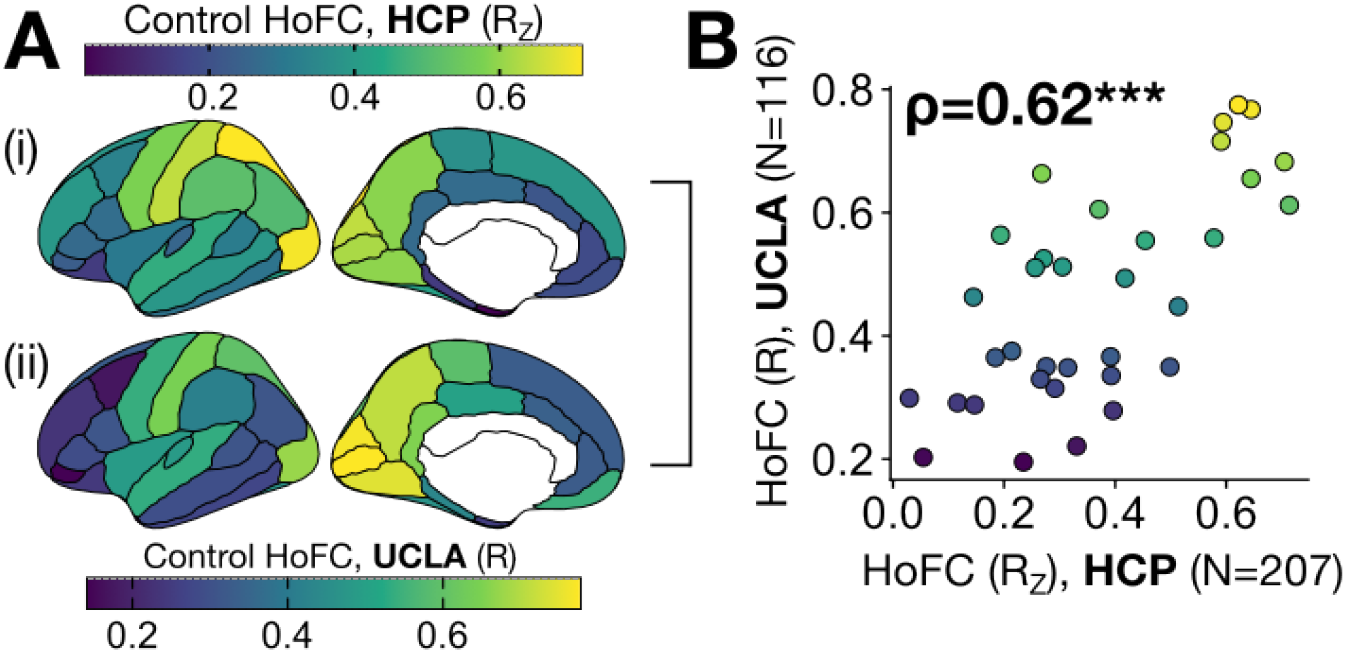
Neurotypical control cohorts show general concordance in homotopic functional connectivity (HoFC) spatial variation. **A**. The regional homotopic functional connectivity (HoFC) values are projected onto the cortical surface for the group averages representing neurotypical control participants from the (i) HCP cohort (N=207) and (ii) UCLA cohort (N=116). Note that the correlation coefficients were *z*-transformed only for the HCP cohort as part of the ENIGMA preprocessing pipeline [48]. **B**. The HoFC values are plotted for each region in the HCP cohort (*x*-axis) and UCLA cohort (*y*-axis), with the Spearman rank correlation (*ρ*) annotated. ^***^, *P <* 0.001, single-hemisphere spin test with 10,000 permutations [56].

**Figure S2.**
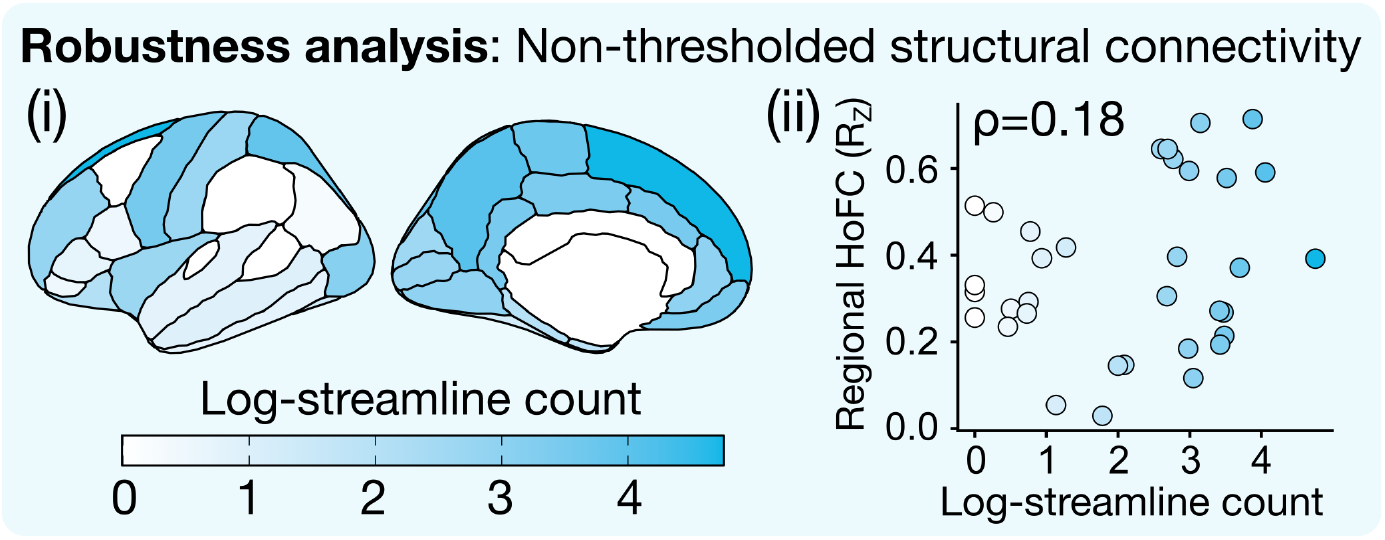
The spatial variation in cortical HoFC is not associated with non-thresholded structural connectivity in a similar HCP cohort. (i) As a robustness analysis, we additionally examined a similar subset of 100 unrelated individuals from the HCP S1200 release as in previous work [82] with a non-thresholded group consensus connectome, derived by computing the mean streamline count per region–region pair across all individuals. Streamline counts are log_10_-transformed; any edges with zero streamlines, or with negative counts after log-transformation, were then set to zero. (ii) The mean log_10_-streamline count connecting each homotopic region pair (*x*-axis) is plotted against the HoFC for each corresponding region (*y*-axis), with the Spearman rank correlation (*ρ*) annotated.

**Figure S3.**
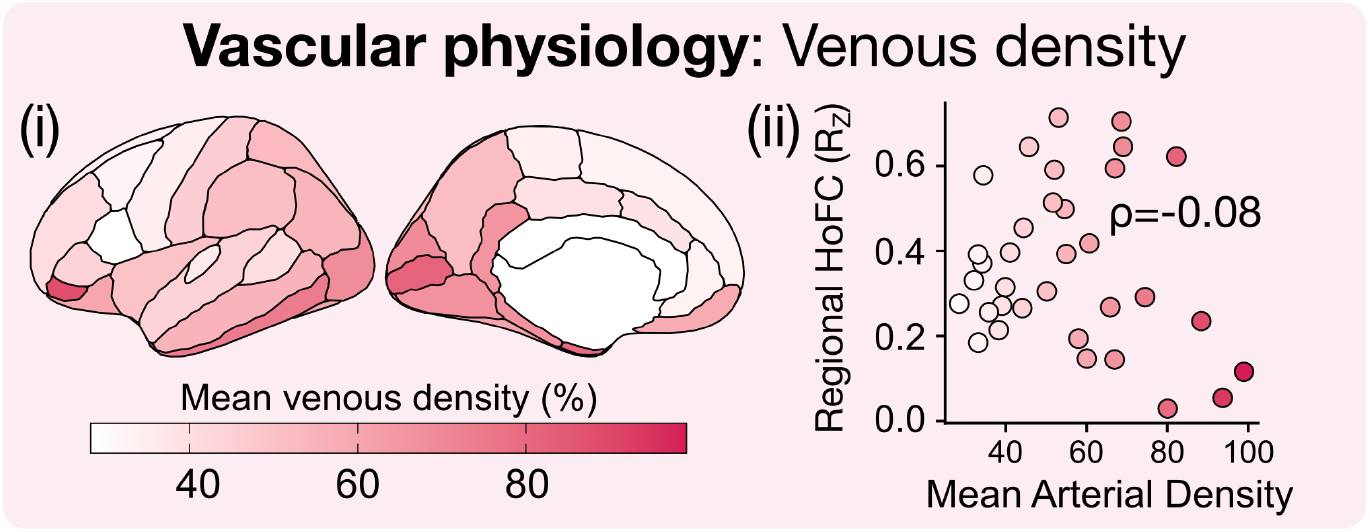
Spatial variation in homotopic functional connectivity (HoFC) is not associated with venous density. (i) The mean venous density in each region is plotted on the cortical surface, averaged between the two hemispheres per region. (ii) The mean hemisphere-averaged arterial density values from the same regions shown in (i) are plotted on the *x*-axis against the HoFC in each corresponding region on the *y*-axis. The Spearman rank correlation (*ρ*) is annotated.

